# Development of host-mimicking legume-based media for robust induction of sporulation in soybean-associated *Cercospora* species

**DOI:** 10.64898/2026.04.10.717671

**Authors:** Nahyun Lee, Juwon Yang, Yoonseo Kwon, Daeyeon Hwang, Jung Wook Yang, Jiyeun Park, Hokyoung Son

## Abstract

*Cercospora* species associated with soybean cause Cercospora leaf spot and purple seed stain, which are major diseases affecting soybean production worldwide and can lead to significant yield and seed quality losses. However, unstable and poor sporulation under laboratory conditions remains a critical challenge, hindering the recovery of genetically homogeneous isolates and the establishment of standardized experimental protocols. These limitations further restrict our understanding of the biology, epidemiology, and pathogenicity of these pathogens. In this study, we developed specialized legume-based culture media derived from soybean and pea tissues to mimic host-associated environmental conditions. We compared the sporulation efficacy of these media with commonly used artificial media, including potato dextrose agar (PDA) and V8 juice agar. Our results demonstrated that legume-based media consistently supported higher levels of sporulation than PDA and V8 across multiple strains, although conidial yields varied depending on the strain and medium concentration. Transcriptional analysis of sporulation-related genes revealed that while *abaA*, *wetA*, and *steA* did not show significant differential expression among media, *velB* exhibited distinct medium-dependent expression patterns. Further evaluation using additional field isolates confirmed that legume-based media provide a more reliable method for inducing sporulation than PDA. Overall, legume-based media represent a practical and effective approach for promoting sporulation in soybean-associated *Cercospora* species under laboratory conditions.

## Introduction

Soybean (*Glycine max* [L.] Merr.) is one of the most economically important crops worldwide, serving as a major source of food, protein, and oil for human and animal consumption, as well as a raw material for diverse industrial applications (Anderson et al., 2019; Hamza et al., 2024). However, global soybean production is frequently constrained by fungal diseases, which cause substantial yield losses and deterioration of seed quality (Soares et al., 2015; Bradley et al., 2021). Among these diseases, *Cercospora*-associated diseases represent a persistent threat to soybean cultivation, leading to foliar symptoms, pod lesions, seed discoloration, and reduced market value (Sautua et al., 2024). In particular, Cercospora leaf spot and purple seed stain occur widely in soybean-growing regions and can result in serious economic damage during severe outbreaks (Li et al., 2019).

Species of the genus *Cercospora* constitute one of the most economically significant groups of fungal pathogens affecting soybean worldwide. Multiple *Cercospora* species have been reported in association with soybean, forming a disease complex that includes leaf spot, pod blotch, and purple seed stain (Soares et al., 2015). Despite their agricultural importance, accurate identification and biological characterization of soybean-associated *Cercospora* species remain challenging, especially at the species level (Hosseini et al., 2023). This difficulty is largely attributable to the high degree of morphological similarity among species and the frequent instability or absence of conidiation under standard laboratory culture conditions, which reduces the reliability of diagnostic morphological characteristics such as conidial shape, size, and septation used for species identification (Souza et al., 2012; Bakhshi et al., 2018; Uppala et al., 2018; da Silva et al., 2025). As a result, routine identification and maintenance of these species are often difficult, severely limiting biological and epidemiological studies.

Accumulating evidence indicates that conidiation and other developmental processes in plant-pathogenic fungi are strongly influenced by host-derived environmental cues (Braunsdorf et al., 2016; van der Does and Rep, 2017). In natural infection sites on leaf and pod surfaces, *Cercospora* species frequently produce abundant conidia, suggesting that specific chemical or physiological signals from host tissues play a key role in regulating reproductive development (Kim et al., 2011; Rangel et al., 2020). These host-associated cues, including phenolic compounds, secondary metabolites, and complex nutrient signals, are largely absent from synthetic media. The pronounced discrepancy between prolific sporulation *in planta* and limited sporulation *in vitro* underscores the importance of culture conditions that more closely mimic the natural host environment (Su et al., 2012). Soybean tissues are rich in phenolic compounds, isoflavones, and diverse carbon and nitrogen sources that may approximate the biochemical environment encountered by pathogens during infection (Albandary Nasser, 2023). Consistent with this, host plant–based media have been widely used to promote sporulation and differentiation in various fungi (Uppala et al., 2018; da Silva et al., 2025). Therefore, this study aimed to develop legume-derived media that mimic host-associated conditions and evaluate their efficacy in inducing and stabilizing sporulation of *Cercospora* species, thereby providing a practical approach for promoting conidiation under laboratory conditions.

## Results

### Development of host-mimicking legume-based media and their efficacy on sporulation

We aimed to establish culture conditions that mimic the host-associated environment to induce conidiation. Thus, we developed two specialized legume-based media, designated as *Glycine max* agar (GA) and *Pisum sativum* agar (PA), prepared using soybean and pea tissues, respectively. To evaluate the influence of host-derived nutrient density on fungal development, both GA and PA were formulated at concentrations of 8% and 16% (w/v). To prepare these media, the respective legume materials were measured according to the desired percentage (8% and 16%), mixed with distilled water, and autoclaved to extract host-associated cues. After cooling, the extracts were squeezed and filtered through sterile gauze, and the final volume was adjusted with distilled water. Agar powder (2%, w/v) was then added, followed by a second autoclaving step to obtain the final solid media (Fig. 1).

**Fig. 1.**
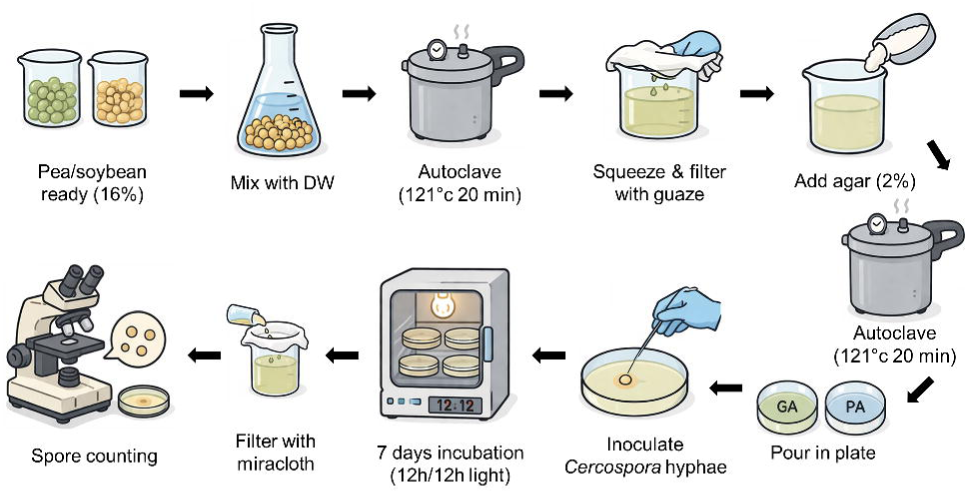
Schematic workflow for conidia induction using legume-based media. Pea or soybean tissues were mixed with distilled water and autoclaved; the extracts were then filtered and supplemented with agar, followed by a second autoclaving step. The media were poured into Petri dishes and inoculated with fresh *Cercospora* hyphae. Plates were incubated under a 12 h light/12 h dark photoperiod. Conidia were harvested by suspending them in distilled water, filtered through Miracloth, and spore concentration was determined using a hemocytometer.

To evaluate the efficacy of these media in promoting sporulation, we tested four *Cercospora sojina* strains originally obtained from soybean in Korea. These strains, designated as 49638, 49639, 49846, and 49847, were acquired from the Korean Agricultural Culture Collection (KACC). Conidial production of these strains was compared across GA, PA, and standard artificial media, including potato dextrose agar (PDA) and V8 juice agar, with water agar (WA) serving as a control. For this evaluation, conidial concentration was measured at 7 and 14 days after inoculation. At 7 days after incubation, strain 49638 failed to produce conidia on PDA, V8 medium, and WA, whereas sporulation was successfully induced on all legume-based media (Fig. 2A). The other three strains, 49639, 49846, and 49847, exhibited similarly limited conidial production on PDA and V8. However, their response to legume-based media varied by strain. For strain 49639, sporulation was markedly enhanced on both GA and PA media, yielding approximately 3.5–4.2-fold and 6.5–10-fold higher spore production than on PDA and V8, respectively. In contrast, while strains 49846 and 49847 also showed improved sporulation on legume-based media, the increase was less dramatic compared to strain 49639. Specifically, for 49846, spore production on 8% GA, 16% GA, and 16% PA was approximately four-fold higher than on V8. Similarly, strain 49847 maintained relatively stable but slightly improved conidiation, with 16% GA showing a significant two-fold increase compared to PDA.

**Fig. 2.**
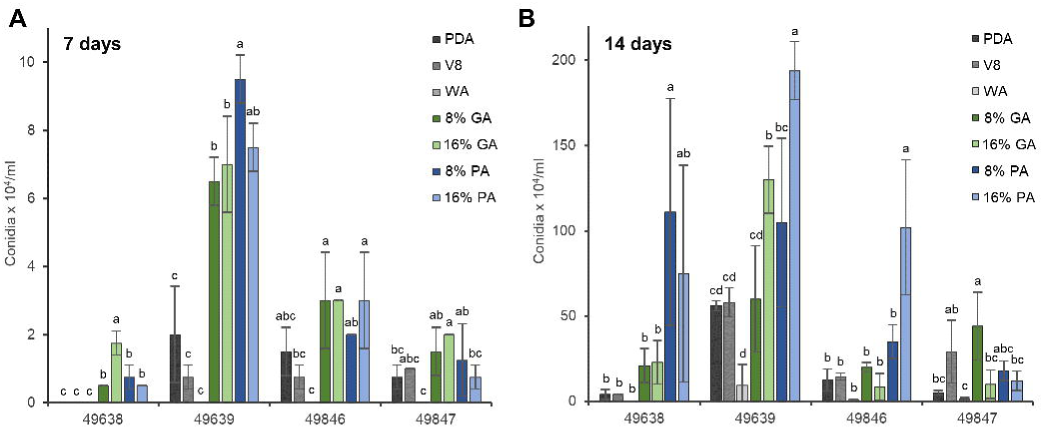
Effects of different media on spore concentration of four *Cercospora* strains. Spore concentrations were measured at (A) 7 days and (B) 14 days after inoculation. Each bar represents the mean spore concentration ± standard deviation (×10 co idia mL ¹). Different letters above the bars indicate significant differences among media based on Fisher’s LSD test (P < 0.05).

After 14 days of incubation, these patterns became more pronounced (Fig. 2B). While strain 49638 produced no spores on PDA and V8 within the first 7 days, conidial induction was observed after 14 days, reaching approximately 4 × 10 conidia mL ¹ on both media. Nevertheless, conidial induction remained stronger on legume-based media compared to these artificial media; yields were approximately 5-fold higher on 8% GA and 16% GA, 28-fold higher on 8% PA, and 18-fold higher on 16% PA relative to both artificial media. Interestingly, there was a notable difference depending on media type. PA media showed a much stronger inductive effect than GA media, with spore production on 8% PA being 5-fold higher than on 8% GA and 16% GA. For strain 49639, while sporulation on 8% GA medium was comparable to that on PDA and V8, spore production was enhanced on other legume-based media. Compared to PDA and V8, conidial yields were approximately 2-fold higher on 16% GA, 1.8-fold higher on 8% PA, and over 3-fold higher on 16% PA. Although no clear difference in induction efficacy was observed based on media type, the concentration of the legume-based media played a significant role, with 16% concentrations inducing significantly more conidia than the 8% concentration.

For strain 49846, no significant enhancement in spore production was observed on most legume-based media, with the exception of 16% PA, which yielded approximately 7- and 8-fold higher conidiation than PDA and V8, respectively. For strain 49847, no substantial enhancement in sporulation was observed on legume-based media compared to the artificial media. Although conidial production on all GA and PA formulations tended to be higher than on PDA, the yields remained comparable to those obtained on V8 agar. Overall, these findings suggest that GA and PA provide a suitable environment for conidiation in *C. sojina*, indicating their potential to induce sporulation across various strains, albeit with varying efficiency depending on the medium concentration and the specific incubation period.

### Optimizing incubation conditions for stable and robust conidial production

To determine the optimal incubation period for sporulation, we monitored conidial production at 3, 5, and 7 days of culture. This time range was selected to identify the earliest time point at which the legume-based media could consistently support robust sporulation across different strains. For this experiment, strains 49638 and 49639 were specifically selected based on their sporulation profiles. Strain 49638 was chosen to represent strains with the lowest baseline sporulation capacity, while strain 49639 showed the most robust inductive response to GA and PA media.

For strain 49638, consistent with previous observations, no conidia were produced on PDA and V8 media throughout the entire 7-day incubation period (Fig. 3A). In contrast, conidiation was initiated on legume-based media starting at 5 days of incubation. Notably, a significant surge in spore production occurred between 5 and 7 days. Across all legume-based media, the conidial yields at 7 days represented a 26- to 52-fold increase compared to the levels recorded at 5 days.

**Fig. 3.**
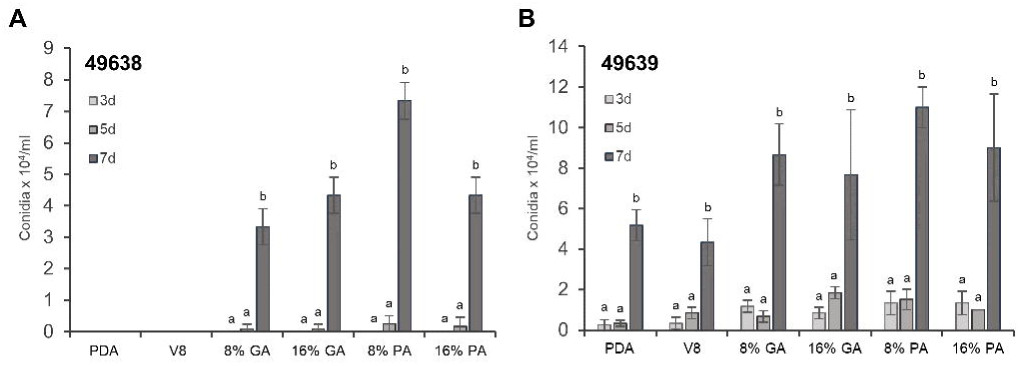
Determination of the optimal time for spore production. Conidia were harvested at 3, 5, and 7 days after inoculation, and their conidial concentrations were measured. Bars represent mean conidial concentrations (×10 conidia mL ¹). Different letters above the bars denote significant differences among media as determined by Tukey’s HSD test (P < 0.05).

A similar pattern was observed for strain 49639 (Fig. 3B). While this strain showed more active sporulation, with conidia detected as early as 3 and 5 days in both legume-based and artificial media, the production levels remained comparable between these two time points. Notably, conidial production increased markedly after 7 days of incubation, significantly outperforming the yields at earlier stages across all tested media.

These results indicate that while the initiation of sporulation is strain-dependent and may occur at earlier stages, substantial and consistent conidial production is most clearly observed across all strains after 7 days of culture. Therefore, a 7-day incubation period is recommended for reliable sporulation induction, ensuring stable yields regardless of the specific strain or culture medium used.

### Relative transcript level of sporulation-related genes

Sporulation in many filamentous fungi is regulated by key developmental regulatory genes, including *abaA, wetA, steA, velB* (Boylan et al., 1987; Breakspear and Momany, 2007; Garzia et al., 2013; Wu et al., 2018). To investigate whether the medium-dependent sporulation pattern in *Cercospora* strains was associated with the differential expression of these regulatory genes, we analyzed their transcriptional levels in strains 49638 and 49639 after 7 days of incubation. Conserved sequences of each target gene were independently identified using genome data from multiple *Cercospora* species available in the NCBI database, and specific primers were designed based on these regions (Table S1).

The resulting expression profiles revealed that while most tested genes lacked a uniform regulatory pattern, *velB* exhibited a transcriptional trend that correlated with the inductive effect of legume-based media across both strains (Fig. 4A). Specifically, in strain 49638, transcript levels of *velB* were consistently higher across all inductive media, showing a more than three-fold increase in GA compared to PDA. Similarly, in strain 49639, significantly elevated transcription was confirmed in 16% GA and 8% PA. In contrast, other regulatory genes did not show a reproducible correlation with the observed sporulation phenotypes under the conditions examined. For instance, *wetA* expression exhibited marginal and inconsistent fluctuation, only showing significant increase in 8% PA for strain 49638 but not in strain 49639 strain (Fig. 4B). Similarly, *abaA* showed no significant difference in expression for strain 49638 and only significantly increased in 16% PA compared to PDA for strain 49639 (Fig. 4C). Furthermore, *steA* showed an increased transcript level only in 8% GA for strain 49638 (Fig. 4D).

**Fig. 4.**
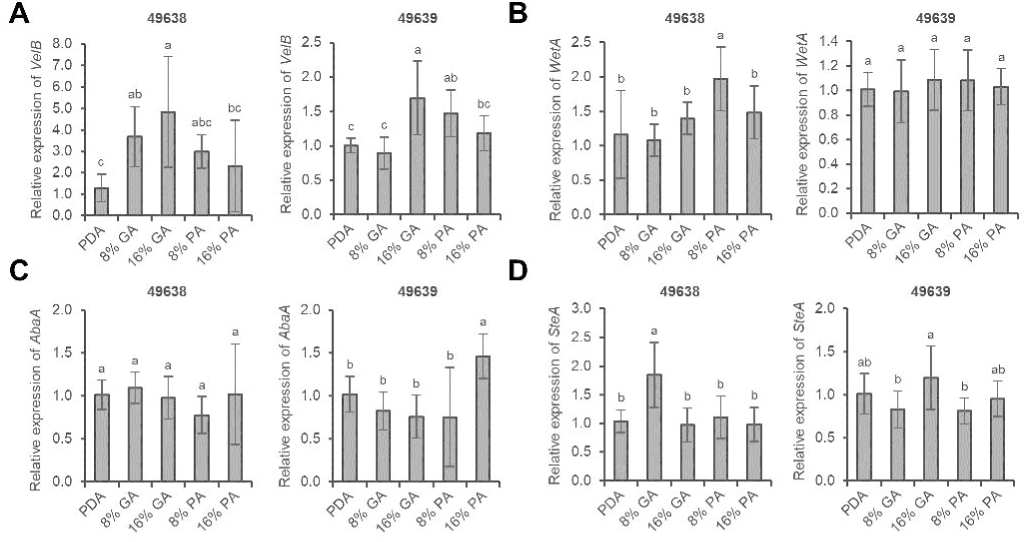
Relative expression of sporulation-related genes in *Cercospora* strains grown on different media. Relative expression levels of VelB (A), WetA (B), AbaA (C), and SteA (D) were measured after 7 days of incubation on PDA, 8% GA, 16% GA, 8% PA, and 16% PA media. Each bar represents the mean ± standard deviation. Bars with different letters indicate significant differences according to Tukey’s HSD test (P < 0.05).

Overall, while a clear linkage between these other regulators and sporulation induction was not confirmed at the 7-day time point, the relatively more consistent induction of *velB* suggests that the velvet complex may be more responsive to medium composition, potentially playing a role in the culture condition–dependent regulation of sporulation in *Cercospora* species.

### Validation of legume-based media using various *Cercospora* field isolates

To evaluate the broader applicability of the developed media, we tested their effectiveness on 16 isolates collected from soybean plants showing purple seed stain symptoms in Korea. These isolates were tentatively identified as *Cercospora* species based on ITS sequence analysis (unpublished data). Conidial production was assessed after 7 days of incubation on PDA, 16% GA, and 16% PA.

While more than half of the tested isolates failed to produce conidia on PDA, every isolate successfully produced conidia on both 16% GA and 16% PA (Fig. 5). Some isolates produced small amounts of conidia on PDA, but their conidial production was consistently higher on 16% GA and 16% PA. For instance, isolate 32 showed markedly increased conidial production on 16% GA and 16% PA compared with PDA, by 27-fold and 11-fold, respectively. Regarding isolate 31, although its conidial production on PDA was comparable to that on legume-based media, it consistently maintained high sporulation rates across all tested media. Altogether, most isolates produced conidia more efficiently on legume-based media than on PDA, suggesting that these media could serve as a new standard for *Cercospora* sporulation.

**Fig. 5.**
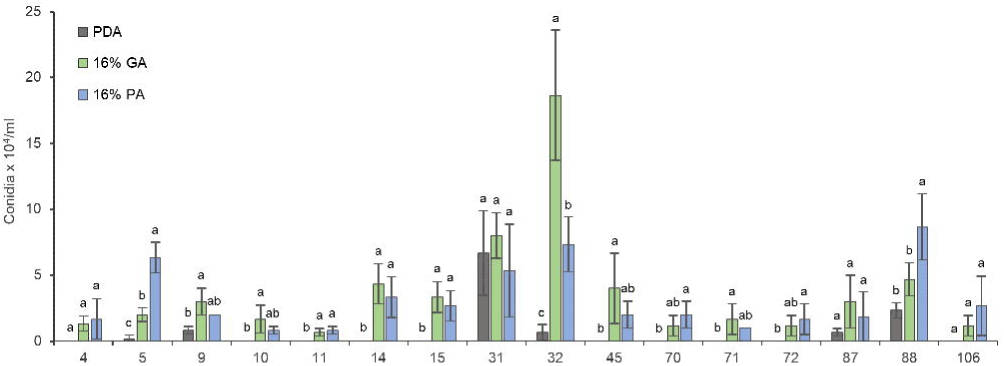
Spore concentrations of field isolates of *Cercospora* species grown on different media. Field isolates obtained from soybean in Korea were cultured on PDA, 16% GA, and 16% PA, and conidial concentrations were measured after 7 days of incubation. The x-axis represents the field isolate numbers used in this study. Bars represent mean ± standard deviation. Different letters indicate significant differences among media according to Fisher’s LSD (P < 0.05).

## Discussion

While numerous studies have examined various media and culture conditions to induce conidiation in *Cercospora* species, establishing a reliable and consistent protocol for sporulation has remained a significant challenge. For instance, during the morphologicial characterization of *Cercospora personata* causing Tikka disease, it was observed that this species did not produce conidia on PDA (Shahdeo Prakriti, 2025). Similarly, although several studies have explored different formulations of V8 agar, PDA, and potato cellulose dextrose agar (PCDA) in attempts to optimize conidial production across multiple *Cercospora* species (da Costa et al., 2020; Sharma et al., 2023; da Silva et al., 2025), sporulation under these conditions is still often inconsistent, necessitating more efficient and standardized methods.

To address these challenges, alternative approaches utilizing host-mimicking media have been introduced as effective strategies. For instance, increased sporulation was observed in *Cercospora lactucae-sativae*, the causal agent of lettuce leaf spot, when media containing higher proportions of host-derived tissues were used instead of PDA (Thomas and Saravanakumar, 2019). Similarly, a pumpkin pathogen *Cercospora citrullina* that failed to produce conidia on PDA produced the highest conidial yields when cultured on media prepared from dried leaves of *Trichosanthes* species (Kathrine Xin Yee Tan, 2020). Furthermore, two cercosporoid species produced abundant conidia when cultured on media supplemented with host leaf decoctions of *Araujia hortorum*, suggesting that host-derived components may play a critical role in sporulation induction (Ramírez et al., 2019). Consistent with these findings, our study verified that legume-based media prepared from soybean and pea tissues not only consistently promoted sporulation in soybean-associated *Cercospora* species but also induced stable conidia production across diverse field isolates. While this approach may be primarily applicable to soybean-associated species, it provides a reproducible and host-relevant platform that overcomes the limitations of conventional media, enabling consistent sporulation and facilitating more reliable morphological and pathological studies.

In achieving effective sporulation, the specific medium formulation is a critical factor and the optimal incubation period plays an equally decisive role. For *Cercospora* species, it has been widely observed that the incubation period required for effective sporulation is not universal. Instead, this timeframe varies significantly depending on both the culture media and the specific species. For instance, while some species, such as *Cercospora* cf*. flagellaris*, can achieve sufficient conidial concentration in as little as 3 days regardless of V8 agar concentration (da Silva et al., 2025), others like *Cercospora medicaginis* have been reported to require prolonged incubation of up to four weeks on PDA and V8 agar media (DjÉBali et al., 2010). These discrepancies highlight the necessity of establishing an optimal incubation timeframe for each specific media system. In this study, even when using two *C. sojina* strains with markedly different sporulation capacities, we observed a consistent and pronounced increase in conidial production after 7 days of incubation on our legume-based media. This reliability was further validated by the fact that conidia were successfully harvested from all 16 tested field isolates, even though their specific species identities have yet to be determined. The consistent performance across these unidentified isolates highlights the broad applicability and robustness of our media system.

To better understand the molecular basis behind this enhanced sporulation efficiency, we examined the expression of key transcription factors involved in the fungal developmental program. The regulatory roles of transcription factors during the sporulation process are well-characterized in many filamentous fungi, including the genus *Aspergillus.* Key transcription factors, such as AbaA, WetA, SteA, and VelB, are strongly expressed during conidiation and function in a stage-specific manner throughout fungal development. In our study, however, most of these genes failed to show a distinct inductive response on the legume-based media, with the exception of *velB,* which exhibited medium-dependent expression patterns (Vallim et al., 2000; Breakspear and Momany, 2007; Tao and Yu, 2011; Wu et al., 2018). This lack of observable induction is likely due to the fact that these genes are regulated in a developmental stage–dependent manner (Breakspear and Momany, 2007). It is highly probable that these regulators are more actively expressed during the early stages of sporulation rather than at the specific sampling time point chosen for this study (Boylan et al., 1987; Tao and Yu, 2011). In contrast to the other tested genes that remained unresponsive, *velB*, a component of the velvet complex, showed consistent induction on legume-based media. This sustained expression suggests that unlike the stage-specific regulators, VelB may function as a persistent regulator during sporulation of *Cercospora* species, as observed in other filamentous fungi. Given that the velvet complex is known to integrate environmental and nutritional cues into the fungal developmental program, the distinct expression profile of *velB* in this study provides evidence for the potential involvement of the velvet complex in the regulation of sporulation in soybean-associated *Cercospora* isolates.

Overall, this study demonstrates that legume-based media not only enhance sporulation efficiency but also provide a reliable and stable method for conidia production in soybean-associated *Cercospora* species. These findings support the utility of host-derived media as an effective alternative to conventional artificial media for inducing sporulation in plant-pathogenic fungi. Furthermore, our results establish a valuable foundation for future investigations into the biology, epidemiology, and disease management of *Cercospora* species associated with soybean.

## Materials and Methods

### Fungal strain and maintenance

Four *Cercospora sojina* strains used in this study were obtained from the Korean Agricultural Culture Collection (KACC). The strains KACC 49846 and KACC 49638 were isolated from Jecheon-si, Chungcheongbuk-do, whereas KACC 49639 and KACC 49847 were isolated from Goesan-gun, Chungcheongbuk-do, and Sunchang-gun, Jeollabuk-do, respectively. These strains were cultured on potato dextrose agar (PDA) for routine maintenance throughout the experiments. For long-term storage, agar blocks containing actively growing mycelia were preserved in 20% glycerol and stored at −80 °C.

### Preparation of media and sporulation induction

To evaluate the effects of medium composition on sporulation, a total of eleven media were used, including PDA, V8 agar, water agar, soybean (*Glycine max*)–based media supplemented at concentrations of 8% and 16%, and pea (*Pisum sativum*)–based media supplemented at concentrations of 8% and 16%. All media were prepared as solid media containing 2% agar and poured into 60-mm Petri dishes (10060; SPL Life Sciences). Fungal strains previously cultured on PDA for 1 week were inoculated onto each medium using 2 × 2 mm agar plugs. The inoculated plates were incubated under fluorescent light with a 12 h light/12 h dark photoperiod. Sporulation was assessed after 7 and 14 days of incubation, and spore formation was examined microscopically. A schematic diagram of the procedure is shown in Fig. 1.

### Quantification of conidia and microscopic analysis

For conidial quantification, aerial mycelia formed on each medium were suspended in 2 mL of distilled water, gently scraped and filtered. The number of conidia in the suspensions was determined using a hemocytometer and expressed as conidia per milliliter.

### RNA isolation and quantitative real-time PCR (qRT-PCR)

After 7 days of incubation, hyphae grown on solid media were harvested by scraping the mycelia without agar and immediately frozen in liquid nitrogen. Total RNA was extracted using the

RNA-spin Total RNA extraction kit (iNtRON, Republic of Korea) according to the manufacturer’s instructions. RNA quality and concentration were assessed using a NanoDrop spectrophotometer (Thermo Fisher Scientific, USA). cDNA was synthesized from the extracted RNA using the SuperScript™III First-Strand Synthesis System kit (Thermo Fisher Scientific). qRT-PCR was performed using iTaq™ Universal SYBR® Green Supermix (Bio-Rad, USA) on a CFX Real-Time PCR System (Bio-Rad), with four biological replicates for each strain and condition.

### Statistical analysis

Experimental results of sporulation are presented as the mean ± standard deviation. Data were analyzed using R version 4.2.3. A one-way analysis of variance (ANOVA) was conducted to evaluate the effect of medium on spore concentration and growth. Following a significant ANOVA result, multiple comparisons were performed using Tukey’s honestly significant difference (HSD) and Fisher’s least significant difference (LSD) test (p < 0.05).

## Acknowledgements

This work was carried out with the support of “Cooperative Research Program for Agriculture Science & Technology Development (No. RS-2024-00397586)”, Rural Development Administration, Republic of Korea.

## Table

Table S1. Primers used in this study

